# AmalgaMo: flexible DNA motif merging

**DOI:** 10.1101/2025.07.29.667561

**Authors:** Orsolya Lapohos, Gregory J. Fonseca

## Abstract

**Motivation:** Inference of candidate upstream regulators via motif enrichment analysis is a common step in the interpretation of genomic data. However, redundancy in motif databases can negatively impact predictive value, especially when relying on regression-based motif enrichment analysis. Although various forms of motif clustering have been used to mitigate problems caused by redundancy, an algorithm optimized for downstream regression-based analysis is needed.

**Results:** We introduce AmalgaMo, an efficient and flexible command line tool for merging highly similar motifs. Using publicly available human datasets, we demonstrate that merging motifs with our optimized settings greatly benefits regression-based motif enrichment analysis and provide detailed documentation that can serve as a reference for researchers inferring upstream regulators from genomic data.

**Availability:** Source code is available on GitHub at https://github.com/lapohosorsolya/AmalgaMo.

## 1 Introduction

Regression-based motif enrichment analysis such as monaLisa Lasso stability selection (Machlab et al., 2022) can provide valuable insight into potential upstream regulators of phenotypes. By modelling changes in epigenomic signals as a function of motif presence with the Lasso penalty, transcription factors (TFs) compete with one another in explaining this variability, somewhat mimicking underlying molecular mechanisms. However, if the regressor matrix contains highly collinear motifs, only one will be selected, ignoring candidates with equal potential.

A seemingly simple solution to this problem is to consolidate highly similar motifs. However, measurement of motif similarity is not trivial, and the optimal level of compression is unknown. Researchers applying motif enrichment analysis expect to obtain a reliable set of candidate upstream regulators that is narrow enough to guide further investigation but broad enough to capture all relevant avenues. Thus, an approach optimized for regression-based enrichment analysis is needed.

Several tools have been developed for grouping similar motifs, such as GMACS (Broin et al., 2015), RSAT matrix-clustering (Castro-Mondragon et al., 2017), GimmeMotifs cluster (Bruse and Heeringen, 2018), abc4pwm (Ali et al., 2022), and universalmotif merge_similar (Tremblay, 2024). Popular motif databases such as JASPAR (Rauluseviciute et al., 2024) and HOCOMOCO (Vorontsov et al., 2024) also have their own reduced motif collections, generated via hierarchical clustering. Despite the number of available tools, a flexible algorithm optimized for a specific down-stream application such as regression-based motif enrichment analysis, does not exist.

Here, we present AmalgaMo, a command line tool that takes as input a set of motifs in any common matrix format and iteratively merges them together using five different parameters that can be tuned for specific applications. We also demonstrate the optimal settings of AmalgaMo for regression-based motif enrichment analysis using paired bulk RNA-seq and ATAC-seq datasets. Further, we leverage the 17,607 available human TF ChIP-seq experiments in the ChIP-atlas (Zou et al., 2022) to thoroughly document the effects of each parameter and guide users tailoring settings to other down-stream applications.

## 2 Methods

### 2.1 Motif representation and metrics

AmalgaMo uses the information-theoretic representation of motifs originally described by Schneider et al. (1986). Briefly, at position *j* of a motif, we denote the Shannon information

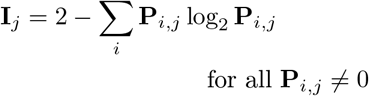

where *i* ∈{A,C,G,T} and **P** is the position-probability matrix (PPM) with rows indexed by *i* and columns indexed by *j*. Then, the total information of a motif is *I* := Σ_*j*_ **I**_*j*_ and we define the *total information ratio* between a pair of motifs *x* and *y*

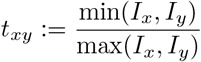

where *I*_*x*_ denotes the total information of motif *x*. Next, we describe the similarity metric used by AmalgaMo: *shared information-weighted cosine similarity*. The shared information of two motifs at position *j* within their alignment region is defined

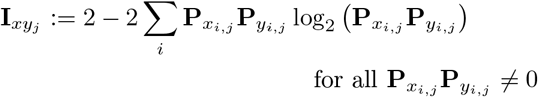

where **P** is padded with background probabilities if there is an overhang. Then, we calculate the cosine similarity at each position along the alignment region and weigh it by the proportion of shared information at that position. Finally, the weighted positional similarities are summed to obtain the scalar similarity score

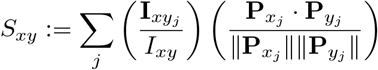

where *I*_*xy*_ is the total shared information between motifs *x* and *y* along their alignment region and 0 ≤ *S*_*xy*_ ≤ 1.

### 2.2 The AmalgaMo algorithm

To ensure that the merged motifs are generated from high-quality matches and to reduce overall compute time, motif pairs are first filtered using three criteria. First, for any pair of motifs (*x, y*), there is a maximum length difference allowance. Second, we set a minimum total information ratio *t*_*xy*_ to consider merging the pair. Third, we find the number of bases within the high-information bounds of both motifs to determine their respective core lengths. These bounds are defined as the first and last positions with at least 1 bit of Shannon information. Then, we apply a maximum core length difference cutoff.

Once a candidate set of motif pairs is obtained, we compute their similarities. We find the best alignment between each motif pair, applying a predefined minimum overlap requirement. Then, the similarity score *S*_*xy*_ is calculated, padding overhangs with background probabilities.

Iterative merging begins with the most similar motif pair. To merge a pair of motifs, their PPMs are simply averaged. If the pair of motifs being merged includes previously merged motifs, each original motif is weighted equally in the final merged PPM. Then, the merged motif is aligned and scored against all other qualifying motifs. This process is repeated until no motif pairs passing the selected similarity score threshold remain.

### 2.3 AmalgaMo parameter selection

To offer flexibility, AmalgaMo has several parameters that can be adjusted by the user. Selecting values for these parameters is relatively simple, as they correspond to intuitive aspects of motif comparison. These include:

- *t*, the minimum total information ratio (default = 0.85);
- *m*, the maximum length difference (default = 2);
- *r*, the maximum core length difference (default = 1);
- *s*, the similarity score cutoff (default = 0.9); and
- *a*, the minimum overlap during alignment (default = 0.9).

For example, a user may want to ensure that certain subtleties are preserved in the merged motifs by using strict settings, such as *m* = 1 and *r* = 0. This parameter set will ensure that for motif sets whose core consensus sequences are almost identical, those with differing positional information contents will be separated (Fig. 1). These settings are explored in the next sections, and detailed in the Supplementary Notes.

**Figure 1:**
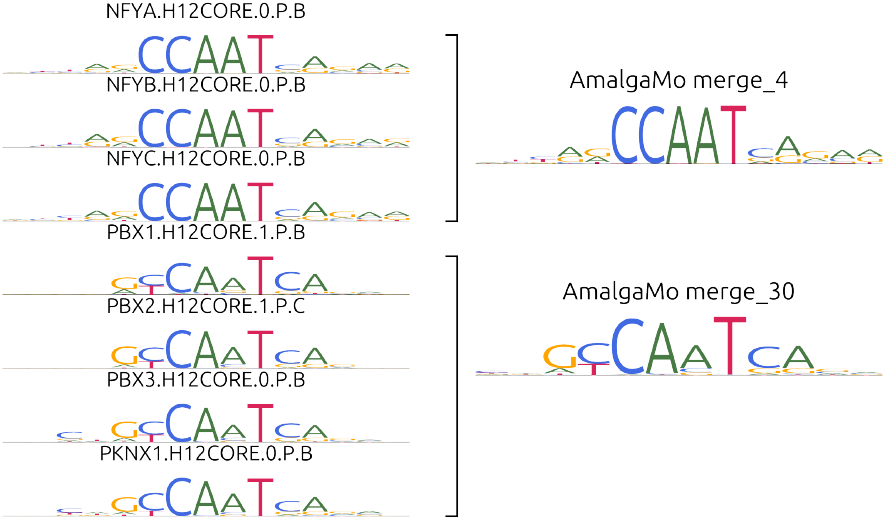
Example of motif merging with relatively strict parameters.

### 2.4 Optimization and evaluation

Although there are many possible use cases for AmalgaMo, we primarily focused on finding the optimal parameter settings for effective regression-based motif enrichment analysis—specifically, the case where the target of the regression is the change in chromatin accessibility between two biological conditions. In order to achieve this goal, we first defined our measures of success. As biological conditions are engendered by differential TF activation, we reasoned that many TFs selected via motif enrichment analysis should be differentially expressed (DE). However, since not all chromatin-bound TFs may be DE, we also considered another measure: a large number of TFs should have detectable mRNA in at least one of the two conditions being compared. Still, it is possible for some relevant TFs to be present in protein form, without any corresponding mRNA. However, since the proteome is context-dependent, we reasoned that if enrichment analysis yields consistent results across different contexts, these metrics reliably capture predictive value.

To calculate these metrics, we needed high quality paired bulk RNA-seq and ATAC-seq data. We obtained such data from three independent sources (Pahl et al., 2024; Li et al., 2019; Mandal et al., 2018) offering biologically homogeneous samples with two comparable conditions (and ≥ 2 replicates per condition). Data processing is described in detail in Supplementary Note 1.

Then, we performed a grid search over three parameters of AmalgaMo (*t, m*, and *r*; keeping *s* = 0.9 and *a* = 0.9), merging motifs from the HOCOMOCO v12 human core motif collection (Vorontsov et al., 2024). We used each of these 27 merged motif sets as input to monaLisa Lasso stability selection (Machlab et al., 2022), supplying the log_2_ fold change (FC) in accessibility between two selected conditions as the target of Lasso regression (details in Supplementary Note 2). Then, we used RNA-seq data to obtain DE TFs for the same two conditions (|log_2_FC| ≥ 1 and FDR *<* 0.05), and counted how many mapped to motifs selected by monaLisa. Considering “positives” to be DE TFs and “negatives” to be non-DE TFs, we calculated Matthew’s correlation coefficient (MCC) for each parameter set. As reference, we also included the original and clustered HOCOMOCO motif sets. Finally, for each dataset, we ranked parameter sets by the number of DE TFs selected, breaking ties using MCC. As the optimal setting, we selected the parameters that consistently ranked first across all datasets.

We also assessed the effects of merging on motif log-odds scores and sensitivity via FIMO using all available ChIP-seq datasets in the ChIP-atlas, as well as motif enrichment analysis with AME (details in Supplementary Notes 3-5).

## 3 Results

### 3.1 Improved predictive value of regression-based motif enrichment analysis

We ran Lasso stability selection from monaLisa (Machlab et al., 2022) using the original and clustered HOCOMOCO v12 motif sets (Vorontsov et al., 2024), and 27 merged motif sets obtained using AmalgaMo with different parameter settings. Using log_2_ FC in accessibility as the target of Lasso regression, the merged motif set generated by AmalgaMo with parameters {*t* = 0.8, *m* = 3, *r* = 1} consistently performed best across all three datasets. This motif set allowed monaLisa to select a greater number of DE TFs, compared to the HOCOMOCO original and clustered motif sets, while reducing the proportion of selected TFs that lacked detectable mRNA (Table 1).

**Table 1:** Evaluation of TFs selected by monaLisa using different motif sets derived from HOCOMOCO, for three datasets. DE: differentially expressed. ND: mRNA not detected. ^∗^ DE TFs defined using only log_2_ FC cutoff due to lack of DE TFs with FDR *<* 0.05.

| dataset | DE TFs | motifs | selected TFs |  |  |
| --- | --- | --- | --- | --- | --- |
|  |  |  | total | DE | ND |
| Pahl<br>2024 | 91 | original | 32 | 10 (31.3%) | 7 (21.9%) |
|  |  | clustered | 97 | 19 (19.6%) | 48 (49.5%) |
|  |  | AmalgaMo | 79 | 20 (25.3%) | 29 (36.7%) |
| Li<br>2019 | 220 | original | 31 | 15 (48.4%) | 13 (41.9%) |
|  |  | clustered | 259 | 71 (27.4%) | 133 (65.5%) |
|  |  | AmalgaMo | 200 | 73 (36.5%) | 126 (63.0%) |
| Mandal<br>2018 | 207* | original | 24 | 2 (8.3%) | 11 (45.8%) |
|  |  | clustered | 77 | 11 (14.3%) | 35 (45.5%) |
|  |  | AmalgaMo | 82 | 16 (19.5%) | 36 (43.9%) |

We provide a more detailed look into motifs selected via AmalgaMo for the dataset from Pahl et al. (2024) in Fig. 2. Here, we can see that, despite keeping the similarity score threshold and minimum overlap constant (*s* = 0.9 and *a* = 0.9) in our grid search, the *t, m*, and *r* parameters had great influence over the motifs selected in the subsequent enrichment analysis (Fig. 2A). This finding highlights the importance of including and tuning such parameters in a motif merging algorithm. Using the optimal AmalgaMo-merged motif set, we found substantial overlap between the selected motifs with the HOCOMOCO original and clustered motif sets (Fig. 2B). However, each one resulted in the selection of unique motifs as a result of differences in grouping and competition enforced by Lasso regression. We also compared the absolute change in gene expression of selected and non-selected TFs. On average, selected TFs had a larger change in gene expression when they were derived from AmalgaMo merged motifs compared to the HOCOMOCO clustered set, but not compared to the original motif set (Fig. 2C). Comparing the union of TFs selected via these three motif sets, there is a clear advantage to grouping motifs for regression-based motif enrichment analysis, especially using AmalgaMo (Fig. 2D). Among AmalgaMo merged hits consisting of more than one motif, 18 DE TFs were identified, 6 of which were also identified via the HOCOMOCO original motif set, and 13 via the HOCOMOCO clustered set (Fig. 2E). Altogether, these results demonstrate that merging motifs using our optimized settings boosts the predictive value of motifs selected via regression-based enrichment analysis.

**Figure 2:**
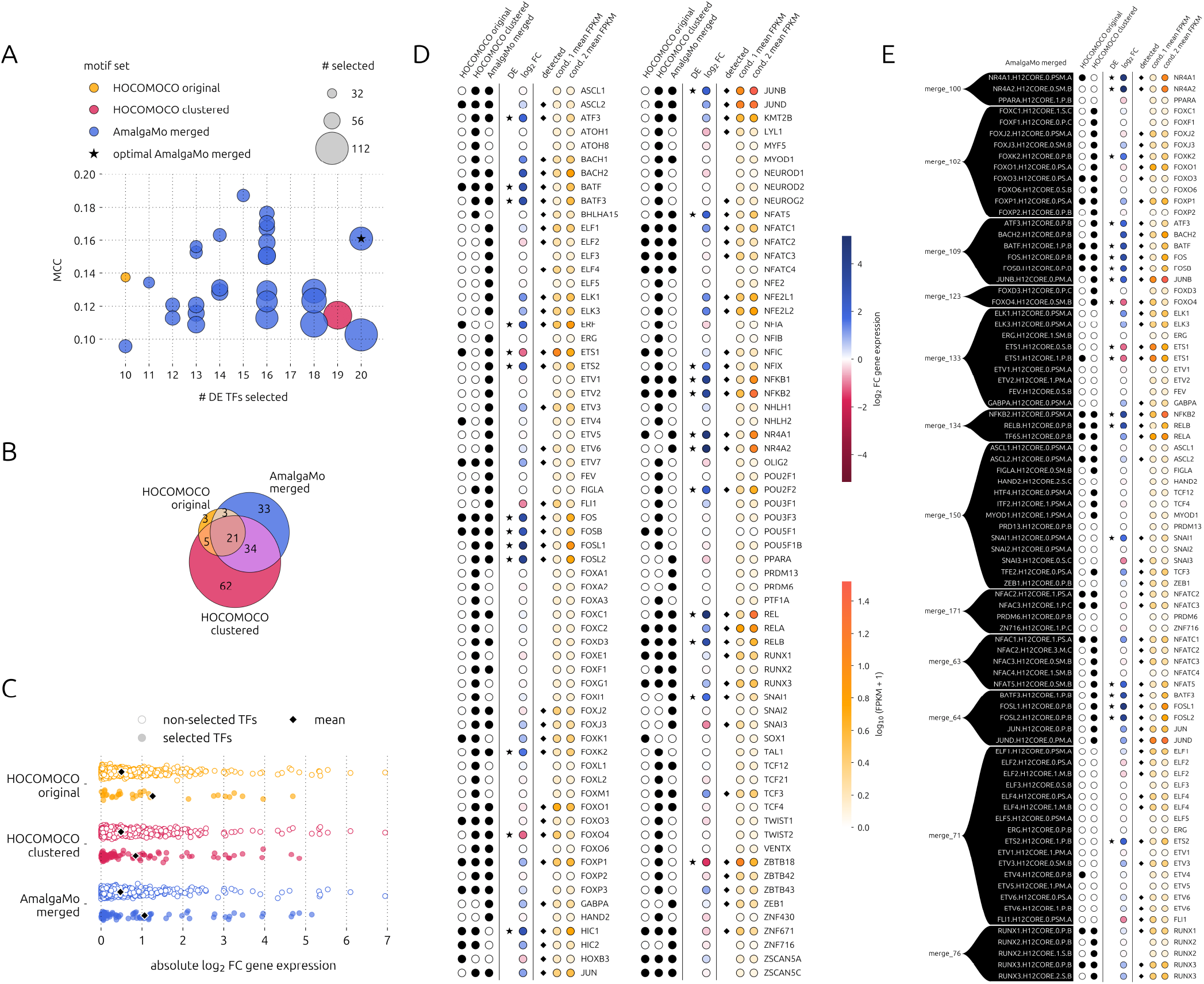
Evaluation of motifs selected via monaLisa Lasso stability selection using a paired RNA-seq and ATAC-seq dataset from Pahl et al. (2024). A. Evaluation metrics for the HOCOMOCO original (orange) and clustered (red) motif sets, as well as AmalgaMo merged motifs (blue) with 27 different parameter sets. Positives are considered to be differentially expressed (DE) TFs. Dot size indicates the total number of selected motifs. The star indicates the optimal AmalgaMo motif set, used in the subsequent figures. B. Venn diagram of the selected motifs. C. Absolute log_2_ fold change (FC) in gene expression of non-selected and selected TFs. Black diamond indicates mean. D. The union of all TFs selected via the three motif sets being compared. Black dots (left) indicate selection. Red and blue coloured dots (center) indicate the log_2_ FC in gene expression of the corresponding TF, with stars marking DE TFs (|log_2_ FC| *>* 1 and false discovery rate *<* 0.05). Yellow and orange coloured dots (right) indicate gene expression (log_10_ FPKM+1), with diamonds marking detection (mean FPKM *≥* 0.5 in at least one condition). E. Same as D, but showing the subset of AmalgaMo merged motifs that were selected, including all corresponding motif versions. MCC: Matthew’s correlation coefficient. FPKM: fragments per kilobase million.

### 3.2 Effect of merging on other aspects of motif enrichment analysis

For a well-rounded evaluation of AmalgaMo, we also assessed changes in motif log-odds scores and sensitivities after merging, using all available human TF ChIP-seq data (17,607 experiments covering 682 TFs) from the ChIP-atlas (Zou et al., 2022). We found that many factors influenced these changes, including motif source data type (ChIP-seq and/or HT-SELEX) and motif quality. Though these findings are not surprising, we provide details and make suggestions for customization of parameter settings in Supplementary Notes 3 and 4.

Finally, although regression-based motif enrichment analysis likely benefits most from merging with Amal-gaMo, we also assessed the results of the independent statistic-based enrichment method AME (McLeay and Bailey, 2010) using merged motifs. We found that, with more relaxed AmalgaMo parameters, the AME ranking of enriched motifs was less consistent with that of the original HOCOMOCO motifs, emphasizing again the importance of tuning these additional constraints. Details are provided in Supplementary Note 5.

## 4 Conclusion

AmalgaMo is a flexible motif merging tool that can be tuned for specific applications. We provide thorough documentation that will allow users to tailor any human database to their needs, or easily produce a non-redundant database for any non-human TF set. We also demonstrate that, with the optimal parameter settings, AmalgaMo can balance the tradeoff between the positive predictive value of motif hits and the number of motifs selected by downstream regression-based motif enrichment analysis. This balance is key to providing researchers with a reliable set of candidate upstream regulators and mitigating misattribution of TF activity.

## 5 Data Availability

Data used to optimize AmalgaMo were downloaded from SRA (accessions: PRJNA960640, PRJNA422918, PRJNA400057).

## Notes

### Competing Interest Statement

The authors have declared no competing interest.

### Funding

- Natural Sciences and Engineering Research Council: 260459

