## Supplementary Material for "AmalgaMo: flexible DNA motif merging"

---

**Supplementary Tables** **15**

**References** **16**

### Supplementary Notes

#### 1 Data processing

Raw FASTQ files were downloaded from three publicly available paired bulk RNA-seq and ATAC-seq datasets on the Sequence Read Archive (accessions in Supplementary Table S1). Reads were mapped to the hg38 reference genome using STAR aligner (Dobin et al., 2013) and then organized with HOMER `makeTagDirectories` (Heinz et al., 2010).

**RNA-seq data.** To obtain normalized expression, we used HOMER `analyzeRepeats` with the `-rpkm`/`-fpkm` option (depending on single-end or paired-end data). For differential expression analysis, we first acquired raw counts using HOMER `analyzeRepeats` with the `-raw` option, and then computed differential expression using HOMER `getDiffExpression` using default settings which rely on DESeq2 (Love et al., 2014).

**ATAC-seq data.** Peaks were called using HOMER `getDifferentialPeaksReplicates` (default: DESeq2) and the `-balanced` and `-all` options, specifying one condition as the target (`-t`) and the other as background (`-b`). The obtained peaks were filtered to those located on autosomes.

#### 2 Lasso stability selection with monaLisa

As the target of monaLisa Lasso stability selection (Machlab et al., 2022), we supplied the $\log_2$ fold change (FC) in chromatin accessibility obtained in the previous step. For constructing the regression matrix, we followed the corresponding Bioconductor vignette, setting the `min.score` parameter of `findMotifHits()` to count motif hits scoring above the $85^{\text{th}}$ percentile, and setting the `cutoff` parameter of `randLassoStabSel()` to 0.8.

#### 3 AmalgaMo parameter effects on motifs

As described in the main text of the article, we merged motifs from the HOCOMOCO v12 human core motif set (Vorontsov et al., 2024), performing a grid search over three parameters of AmalgaMo: the minimum total information ratio $t \in \{0.80, 0.85, 0.90\}$, the maximum length difference $m \in \{1, 2, 3\}$, and the maximum core length difference $r \in \{0, 1, 2\}$. Since we were interested in evaluating the effects of these additional constraints on the resulting merged motifs, we kept the similarity score cutoff and minimum alignment overlap parameters constant ($s = 0.9$ and $a = 0.9$, respectively).

##### 3.1 Basic evaluation of AmalgaMo merged motifs

On first examination, we observed that relaxing the constraints imposed by the parameters tested in our grid search lead to a greater reduction in the size of the merged motif set generated by AmalgaMo (Supplementary Figure SN3.1A). The greatest reduction achieved was 51% with the least strict parameter set, and the least reduction achieved was 26% with the most strict parameter set.

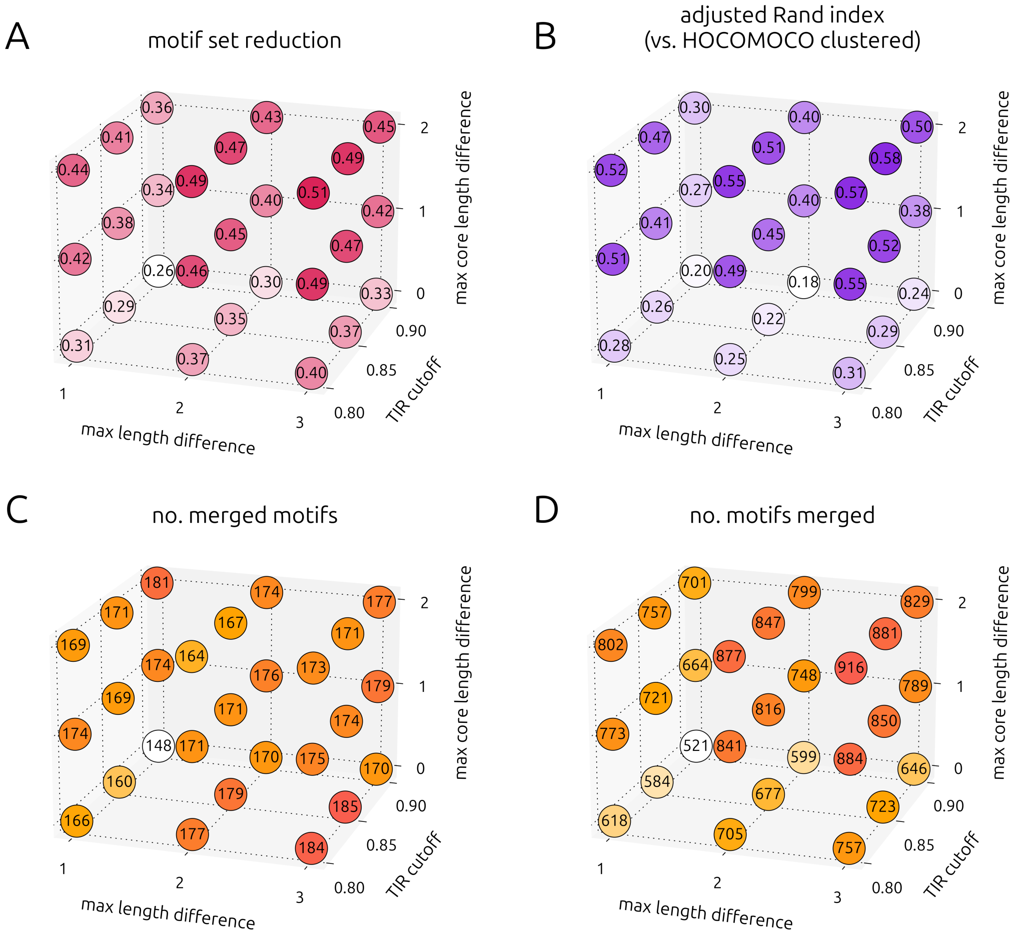

**Supplementary Figure SN3.1:** Basic metrics of motif sets obtained via AmalgaMo.

To compare these merged motif sets to the HOCOMOCO v12 clustered motif set, we computed the Rand index (RI) and adjusted Rand index (ARI) using `rand_score()` and `adjusted_rand_score()` from scikit-learn (Pedregosa et al., 2011). Although motifs were grouped very similarly across all comparisons (RI $> 0.98$), HOCOMOCO clusters tended to be coarser. As a result, we examined differences in granularity using ARI, which varied between 0.18 and 0.58 (Supplementary Figure SN3.1B). Unsurprisingly, the pattern in ARI was generally consistent with the degree of motif set size reduction by AmalgaMo.

Next, we looked at the number of merged motifs; i.e., the number of motifs output by AmalgaMo that were derived from more than one input motif. This value ranged from 148 to 185, with the greatest number given by the parameters $\{t = 0.85,\ m = 3,\ r = 0\}$ (Supplementary Figure SN3.1C). However, the number of motifs merged (input motifs that were combined with others) exhibited a different trend, with this parameter set falling on the lower end of the spectrum (Supplementary Figure SN3.1D). Together, these metrics indicate that with more relaxed (higher values of) $t$ and $m$, restricting the maximum core length difference ($r$) results in a more fine-grained merged motif set, ensuring the formation of a greater number of merged motifs using fewer input motifs.

##### 3.2 Normalized motif edit distance

To independently compare merged motifs with their original components, we devised an alternative strategy that relies on base-wise log-likelihoods at each position. First, the original motif is aligned to the merged motif, padding overhangs. Next, position-weight matrices (PWMs) are calculated for both motifs. The weight of a base $i \in \{A, C, G, T\}$ at position $j$ is given by

$$\mathbf{W}_{i,j} = \log_2 \frac{\mathbf{P}_{i,j}}{\mathbf{b}_i}$$

where $\mathbf{b}$ is the vector of background probabilities for each DNA base (here, $\mathbf{b}_{ACGT} = [0.275, 0.225, 0.225, 0.275]$). Then, a binary matrix $\mathbf{U}$ is calculated for each motif by element-wise thresholding of the PWM:

$$\mathbf{U}_{i,j} = \begin{cases} 1 & \mathbf{W}_{i,j} > 0 \\ 0 & \mathbf{W}_{i,j} \leq 0 \end{cases}$$

such that $\mathbf{U}$ indicates the bases that have a greater-than-random likelihood of occurrence. Finally, we define a new metric called *normalized motif edit distance* (NMED), which is related to string and graph edit distance metrics. The NMED between two motifs is defined as the minimum number of modifications needed to transform the binarized original PWM ($\mathbf{U}_x$) into the binarized merged PWM ($\mathbf{U}_y$), normalized to motif length, $\ell$. Formally,

$$\text{NMED}_{xy} = \frac{1}{4\ell} \sum_i \sum_j |\mathbf{U}_x - \mathbf{U}_y|_{i,j}$$

such that $0 \leq \text{NMED} \leq 1$.

We assessed NMED of merged motifs with respect to their original components (Supplementary Figure SN3.2) and found two consistent trends. First, the mean NMED across all merged motifs has an inverse relationship with the total information ratio cutoff ($t$). Second, the mean NMED increases with the maximum core length difference ($r$). There is no clear pattern when varying maximum length difference ($m$), but the smallest NMED was obtained using the smallest length difference allowance. Thus, it is especially important to consider the $t$ and $r$ parameters in addition to a similarity score threshold when merging motifs.

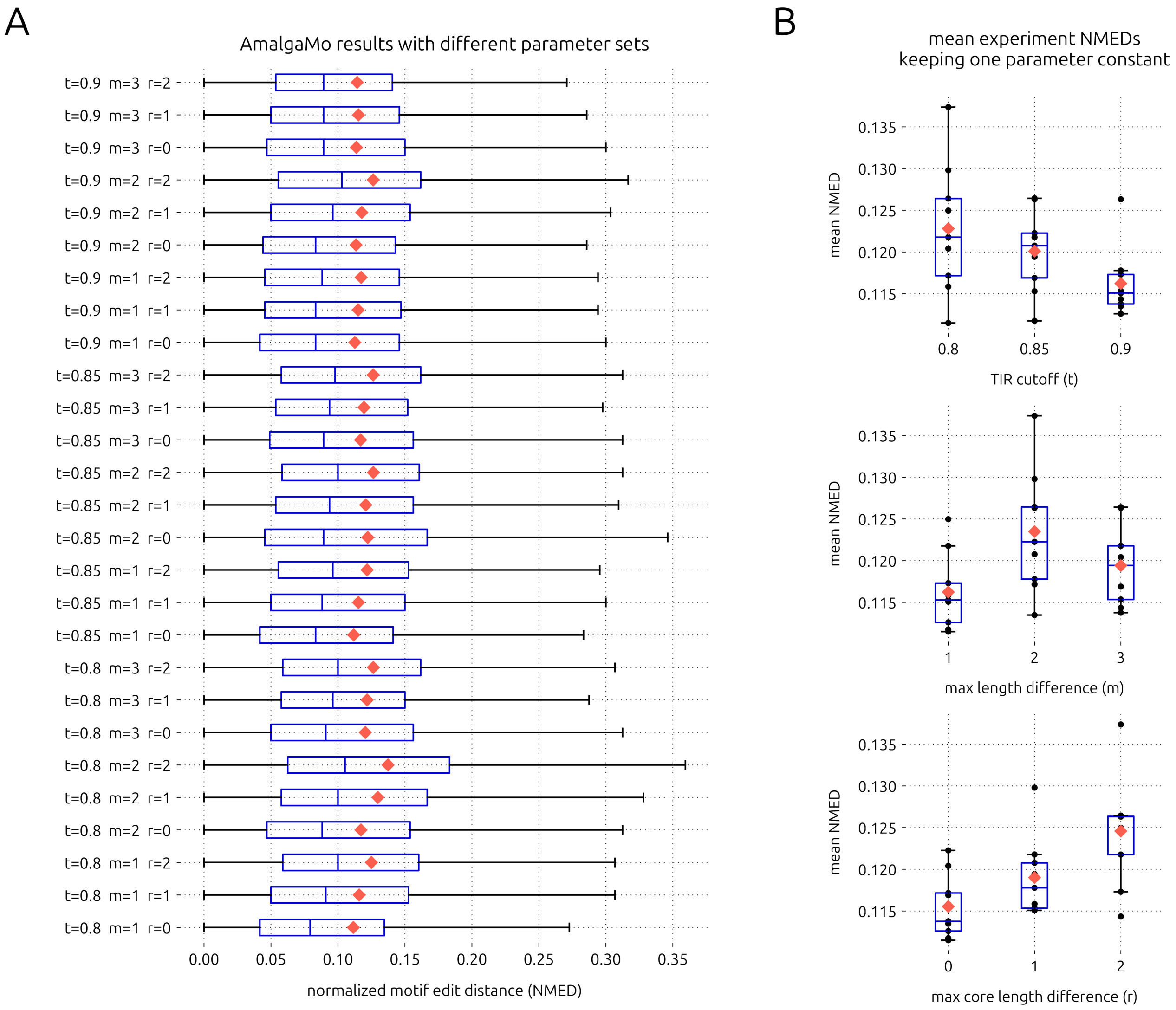

**Supplementary Figure SN3.2:** NMED of AmalgaMo-merged motifs versus their original components using different parameter sets. A. Boxplots showing results from every parameter set. Red diamonds indicate means. B. Boxplots showing mean NMED from each experiment in A, keeping one parameter constant. TIR: total information ratio.

##### 3.3 Motif scores and sensitivity

**Evaluation criteria.** To test the merged motifs against experimental data, we downloaded all high quality human TF ChIP-seq peaks ($q$-value $< 10^{-50}$) aligned to the hg38 reference genome from the ChIP-atlas (Zou et al., 2022). Then, for a set of merged motifs, we took the subset with available ChIP-seq data for at least 3/4 of the corresponding original motifs in order to ensure a fair evaluation. While the ChIP-atlas has accumulated a large number of ChIP-seq experiments over the years, there is still a substantial amount of missing data. Thus, we were able to evaluate only 40-47% of our merged motifs, corresponding to 32-35% of all original motifs that were merged (Supplementary Figure SN3.3). We note that, although there was no apparent trend in the number or proportion of testable motifs by parameter, the subsequent motif evaluation results should be considered carefully and in combination with the NMED results, given the number of motifs that could not be evaluated empirically.

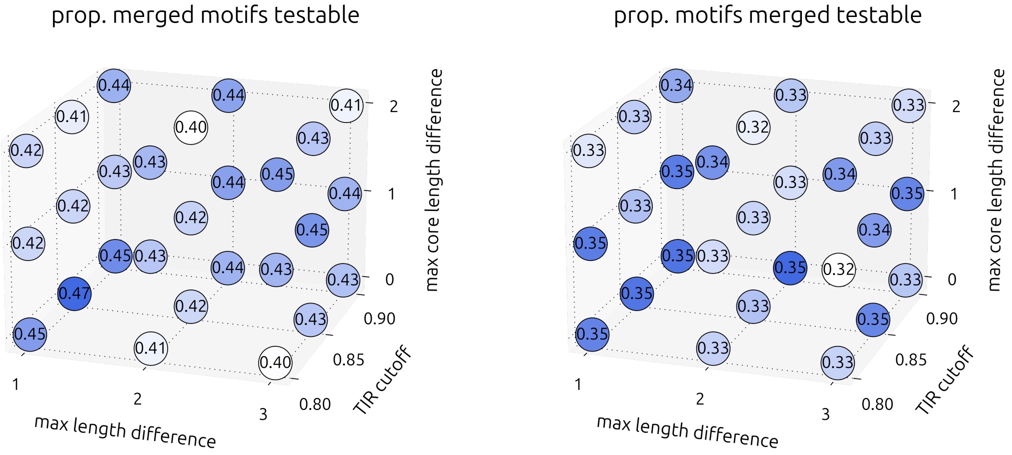

**Supplementary Figure SN3.3:** Proportion of testable motifs in each AmalgaMo merged motif set. Merged motifs testable: output motifs derived from more than one input motif, with high quality data available in the ChIP-atlas for at least 3/4 of the input motifs. Motifs merged testable: original component motifs (inputs) that were merged such that the minimum ChIP-atlas data criterion was met.

**Log-odds scores.** For all ChIP-seq peaks along chromosome 6, we computed log-odds scores for corresponding original and merged motifs at every position, and took the maximum within each peak. The log-odds score of a query sequence $q$ is given by

$$\sum_i \sum_j \mathbf{W}_{ij} \cdot \mathbf{1}_i(q_j)$$

where $\mathbf{1}_i$ is the indicator function which outputs 1 if $q_j = i$ and 0 if $q_j \neq i$. Finally, we calculated the coefficient of determination ($R^2$) between the maximum log-odds scores for each ChIP-seq dataset corresponding to each original motif and its associated merged motif. We also counted the frequency of improvement in log-odds score from the original to the merged motif as a complementary measure of success. These metrics were then averaged across datasets for each original motif. We opted to calculate the median (rather than the mean) in order to reduce the influence of outliers.

**Sensitivity.** As another measure of comparison between the original and merged motifs, we used the MEME Suite's `FIMO` (Grant et al., 2011) to scan chromosome 6 for individual motif occurrences and compared FIMO hits with $p$-value $< 10^{-4}$ to corresponding ChIP-seq peaks from the ChIP-atlas. We considered FIMO hits coinciding with ChIP-seq peaks as true positives and ChIP-seq peaks without FIMO hits as false negatives. From these values, we calculated sensitivities for all available ChIP-seq datasets and their corresponding original and merged motifs. Then, we calculated the median fold change in sensitivity across datasets to systematically compare the original and merged motifs.

**Example.** First, we demonstrate the metrics described in the two paragraphs above, using the merged RUNX family motifs in Supplementary Figure SN3.4. In general, the maximum log-odds scores are very similar between the original and merged motifs. Three of the five motifs (RUNX1.H12CORE.0.P.B, RUNX2.H12CORE.0.P.B, and RUNX3.H12CORE.0.P.B) have high $R^2$ across all datasets, with minimal improvement in scores. However, two motifs (RUNX2.H12CORE.1.S.B and RUNX3.H12CORE.2.S.B) have low $R^2$ with substantial improvement in log-odds scores after merging. These changes are also reflected in the merged versus original sensitivities, where we can see that the original motifs' sensitivities are maintained for the most part, and even improved upon in some cases. The reader may notice that improvement within this example coincides with "S" category motifs—this relationship is explored systematically in Supplementary Note 4.1.

**Overall results.** Given that the act of merging motifs eliminates small differences when averaging motif matrices, we expected the changes in sensitivity to be affected by the degree of parameter strictness. We found several notable trends when examining individual parameter effects (Supplementary Figure SN3.5). Interestingly, the median improvement in log-odds scores within ChIP-seq peaks increased with the maximum length difference allowance. The same trend was observed for median fold change in sensitivity. The strongest trend we observed was the decrease in median fold change in sensitivity with respect to the maximum core length difference parameter. With this parameter set to 0, the (testable) merged motifs were guaranteed to maintain their sensitivities compared to their original components, regardless of the other parameters. However, we note that the worst-case scenario in terms of median sensitivity loss within our grid search was a decrease of only 1.2%, which we deemed acceptable.

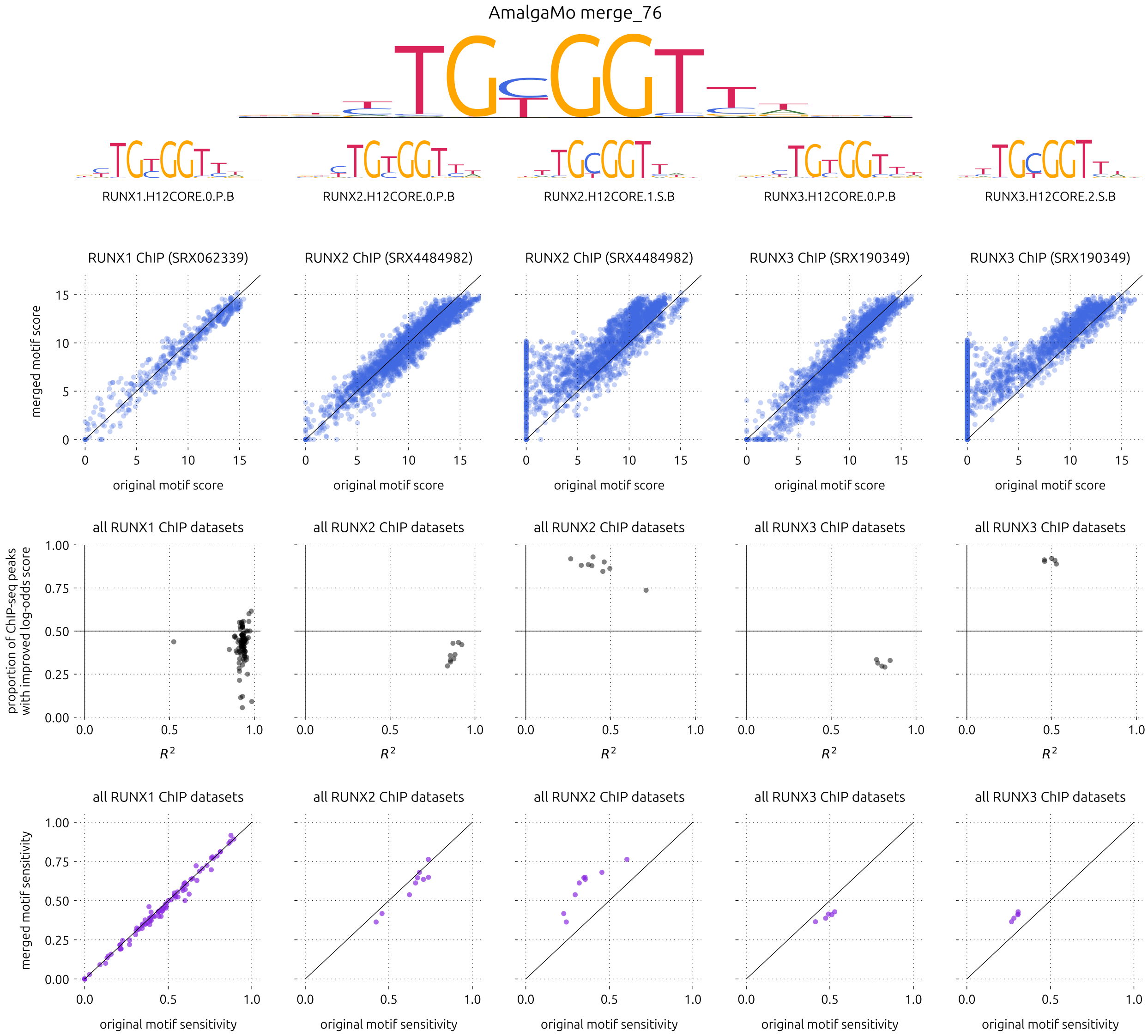

**Supplementary Figure SN3.4:** Changes in motif scoring and sensitivity when merging RUNX family motifs. The large logo shows the merged motif, which is composed of the five motifs underneath. The top row of scatter plots (blue) show the maximum motif scores within ChIP-seq peaks from randomly selected corresponding datasets. The second row of scatter plots (black) show the proportion of ChIP-seq peaks with improved log-odds score versus the coefficient of determination of merged and original motif log-odds scores, for all available ChIP-seq datasets in the ChIP-atlas.

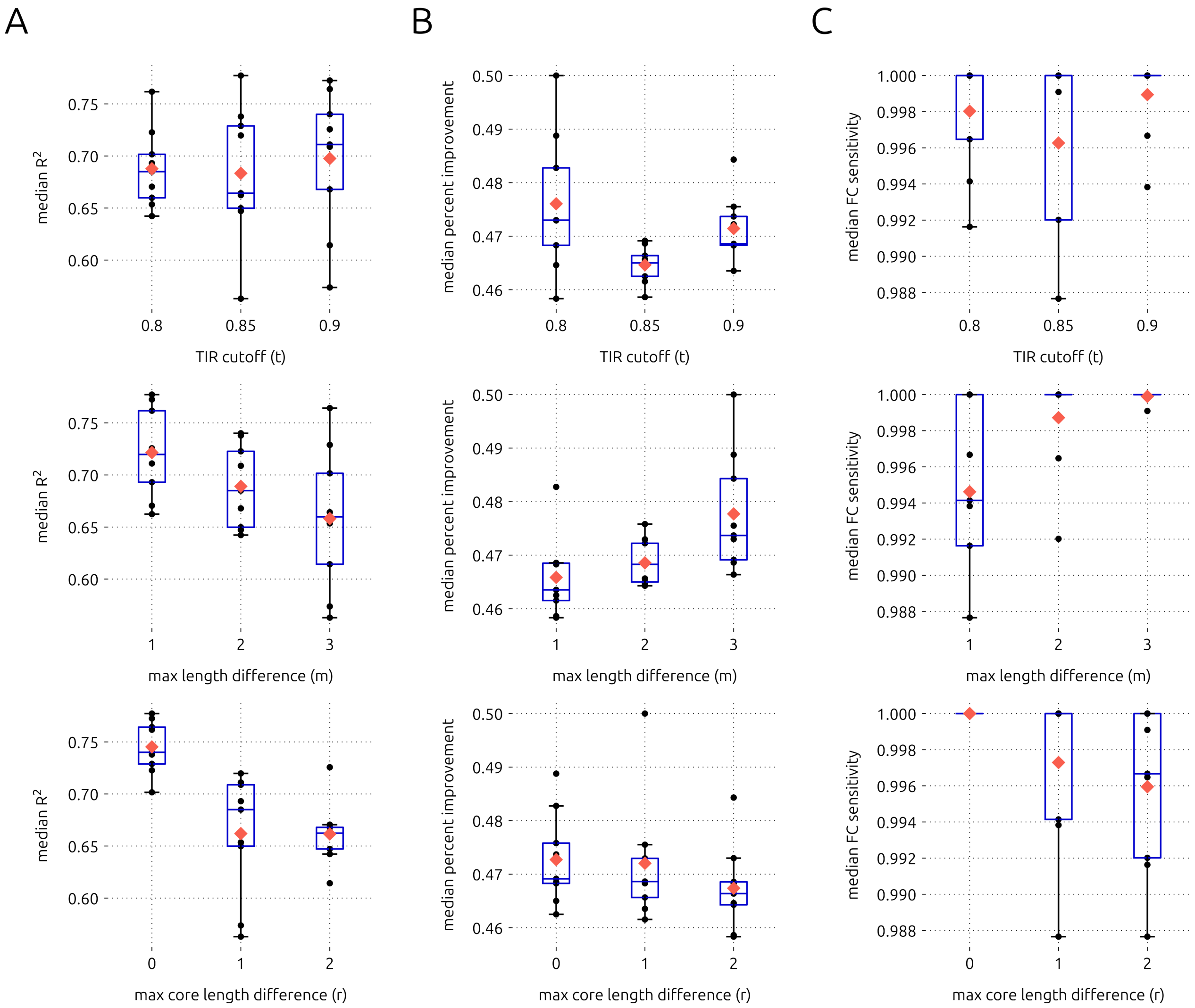

**Supplementary Figure SN3.5:** AmalgaMo parameter effects on motif scores and sensitivities. A. Median $R^2$ of maximum log-odds scores of original versus merged motifs in all available corresponding ChIP-seq datasets. B. Same as A, but showing median proportion of ChIP-seq peaks with improved max log-odds scores. C. Median fold change (merged/original) in motif sensitivity within ChIP-seq peaks as detected by FIMO.

### 4 Additional considerations for AmalgaMo parameter selection

#### 4.1 Motif source data

Although it is widely known that (Methyl-)HT-SELEX and ChIP-seq based motifs differ, we wanted to determine how motif source data impact merged motifs. The HOCOMOCO database (Vorontsov et al., 2024) labels each motif as follows:

**P:** motif derived from ChIP-seq data

**S:** motif derived from HT-SELEX data

**M:** motif derived from Methyl-HT-SELEX data

We compared the motif sets obtained using the least and most strict parameter sets in terms of maximum log-odds scores and sensitivities within ChIP-seq peaks (Supplementary Figure SN4.6). We found that, in both cases, motifs derived from only (Methyl-)HT-SELEX data tended to be most affected by merging. On average, these were the motifs that benefited most, while those derived from ChIP-seq data often did not benefit. This result makes sense, given that the evaluation metrics rely on ChIP-seq data. Further, a small number of input motifs lost a great deal of sensitivity after being merged (though most of these motifs had very few supporting datasets), but constraining AmalgaMo parameters reduced these losses. These findings suggest that care should be taken with regards to the source data of motifs input to merging algorithms, and when selecting parameters for AmalgaMo. Depending on the downstream application, it may also be preferable to exclude certain motifs or even an entire category of motifs.

#### 4.2 Motif quality

Next, we explored the relationship between motif quality and merged motif scoring and sensitivity. Conveniently, the HOCOMOCO database (Vorontsov et al., 2024) also labels each motif with a quality rating on the ABCD scale:

**A:** motif derived using both (Methyl-)HT-SELEX and ChIP-seq data

**B:** motif found in at least two experiments of the same type

**C:** motif passed expert curation but found in only one experiment

**D:** "legacy" motif, not benchmarked

Dividing the motifs merged by AmalgaMo into A/B and C/D quality groups, we found that the number of C/D motifs merged more than doubled from the most strict to the most relaxed parameter setting in our grid search (from 73 to 153). While this increase is not cause for concern, users should keep it in mind when selecting the input motif set and AmalgaMo parameters.

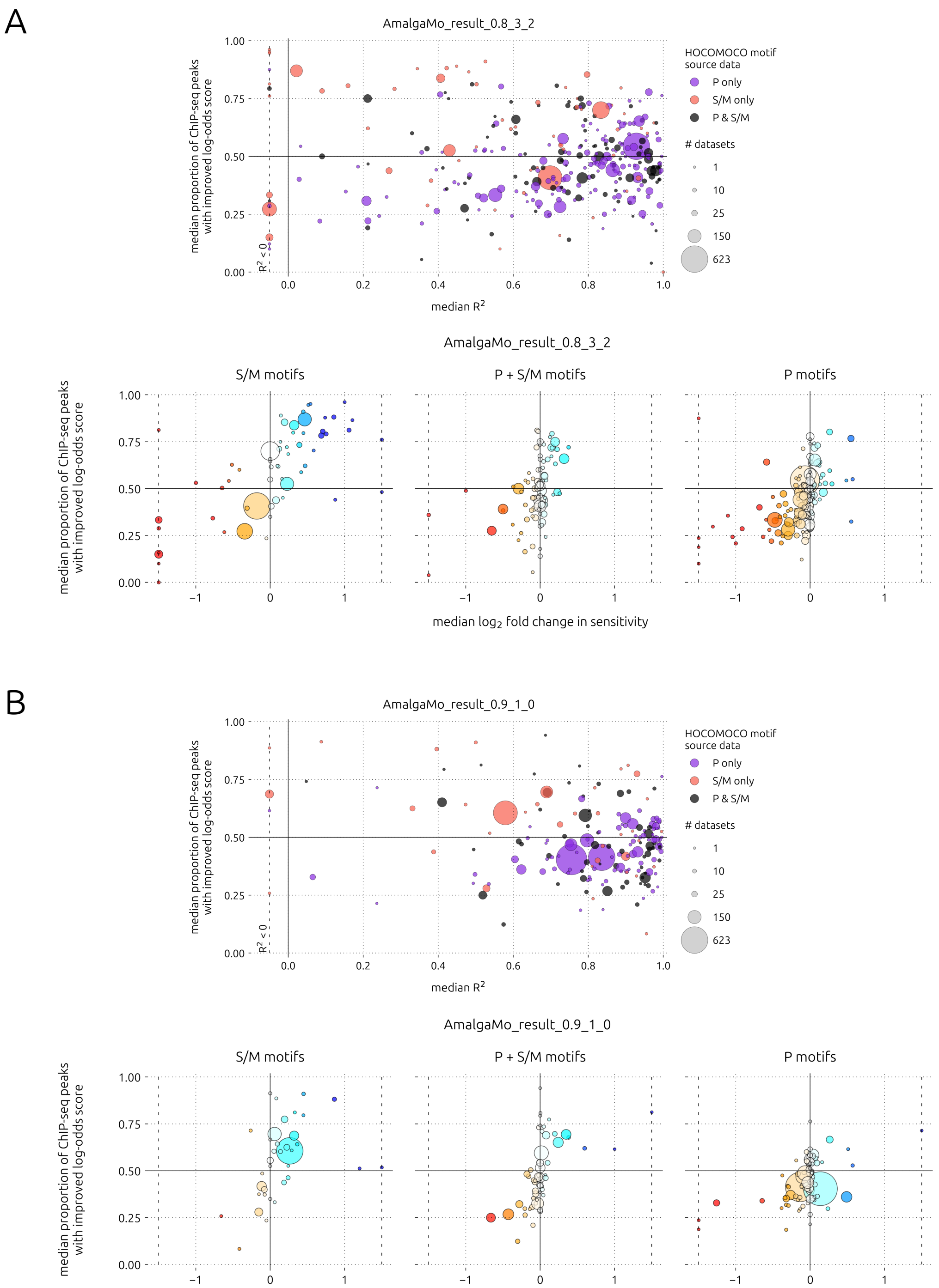

**Supplementary Figure SN4.6:** Empirical evaluation of motifs merged, by motif source data. A. Motif set obtained using the most relaxed parameter set. B. Motif set obtained using the most strict parameter set. P: ChIP-seq. S: SELEX. M: methyl-SELEX. Marker area is proportional to the number of datasets used for evaluation. Markers on the dashed vertical lines indicate values outside the range of the $x$-axes.

Among those motifs merged that could be evaluated with ChIP-seq data, quite a few A/B quality motifs experienced a decrease in sensitivity after merging using relaxed parameters (Supplementary Figure SN4.7). This number decreased substantially when using the strict parameter set. We also found that, regardless of parameter set, the proportion of C/D quality motifs with extreme changes in sensitivity was greater compared to A/B quality motifs. Thus, for downstream applications calling for high confidence, users may want to exclude C/D quality motifs.

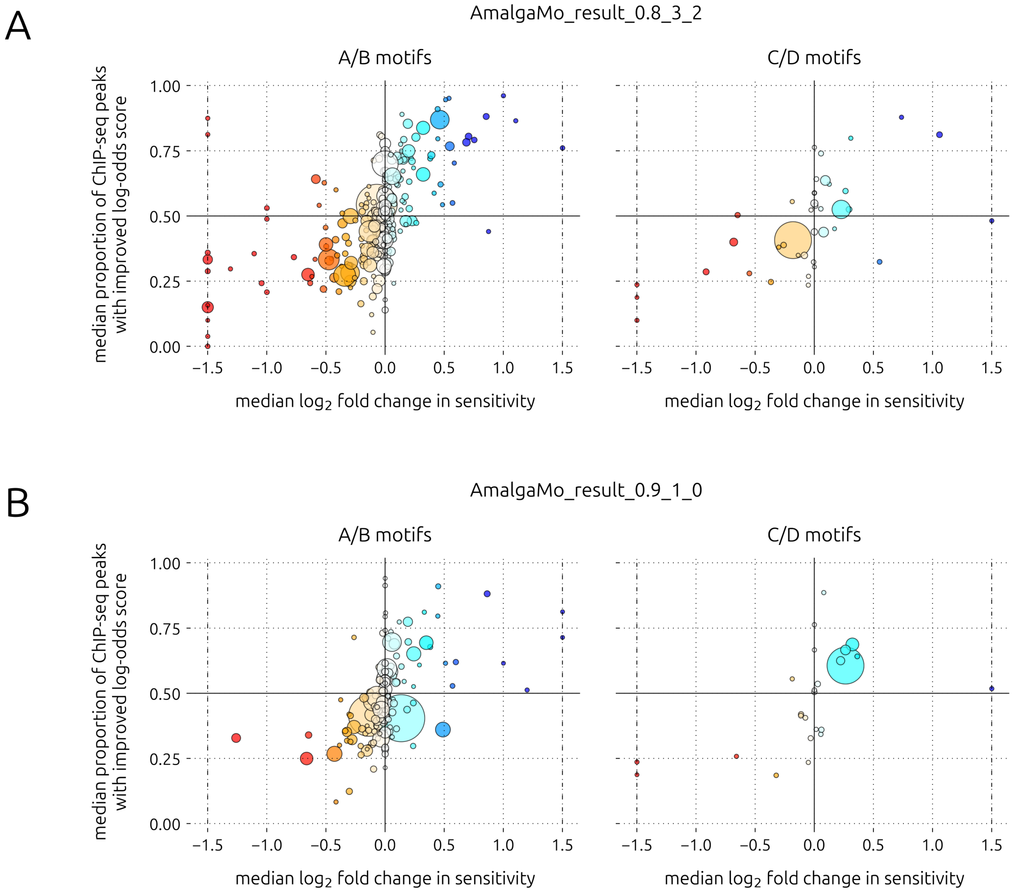

**Supplementary Figure SN4.7:** Empirical evaluation of motifs merged, by motif quality. A. Motif set obtained using the most relaxed parameter set. B. Motif set obtained using the most strict parameter set. Marker area is proportional to the number of datasets used for evaluation. Markers on the dashed vertical lines indicate values outside the range of the $x$-axes.

#### 4.3 Motif database

Another consideration worth mentioning has to do with the different *de novo* motif finding algorithms and curation strategies employed by motif databases. These aspects fall outside the scope of this study, but we wanted to bring users' attention to key differences that are likely to impact the ideal parameters for merging motifs. We compared human motif collections from three human databases: JASPAR (Rauluseviciute et al., 2024), HOCOMOCO (Vorontsov et al., 2024), and CIS-BP (Weirauch et al., 2014).

We found that these collections differed considerably in terms of motif characteristics and TF redundancy (Supplementary Figure SN4.8). JASPAR has the fewest human motifs (720), and they are shorter on average, with greater information content per position, and shorter tails (regions flanking the "core", with positional information content less than 1 bit). These properties make JASPAR motifs seem more exact, with base occurrence probabilities diverging eminently from a uniform distribution. This collection also has relatively low TF name redundancy. On the other hand, HOCOMOCO has approximately twice the number motifs in JASPAR (1443). HOCOMOCO motifs also have longer tails, increasing the average motif length. In general, these motifs have less information per position, but this difference is minute when looking only at the high-information region (motif core). HOCOMOCO also has slightly more redundancy in terms of TF name frequency, due to the database storing multiple subtypes of motifs per TF when empirical data supports their existence. Finally, CIS-BP is a comprehensive database, having 6607 human motifs at the time of this analysis, housing collections from many other databases (including JASPAR and HOCOMOCO) as well as inferred motifs. Unlike JASPAR and HOCOMOCO, CIS-BP is not curated and therefore exhibits high redundancy. In terms of motif length and tail size, CIS-BP falls in between JASPAR and HOCOMOCO.

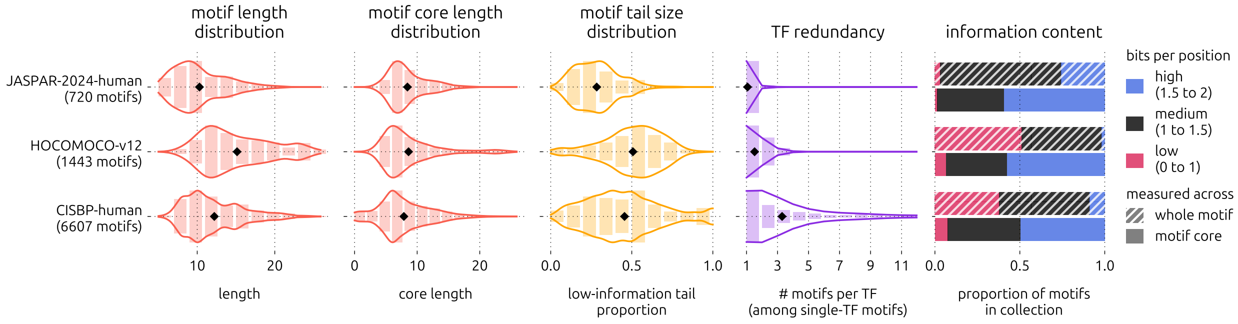

**Supplementary Figure SN4.8:** Comparison of human motif characteristics between three popular databases.

For AmalgaMo users, we have two suggestions based on input motif set characteristics. First, for motif collections with longer average tail sizes (such as HOCOMOCO), the user should consider relaxing the maximum length difference parameter ($m$). Second, for motif collections with a very low proportion of low average positional information content motifs (such as JASPAR), the user should consider setting the maximum core length difference parameter ($r$) to 0.

#### 5 Motif enrichment analysis with merged motifs

To evaluate the change in ranking of enriched motifs by an independent statistic-based method, we ran the MEME Suite's `AME` (McLeay and Bailey, 2010) on the dataset from Pahl et al. (2024). We supplied peaks from one condition (stimulated) as query sequences, and peaks from the other (unstimulated) as control sequences using the `--control` option.

We found that some motifs were promoted in rank by association with highly enriched motifs, while others were bumped down the list in significance (Supplementary Figure SN5.9). Since AME tests each motif independently, these changes can be attributed entirely to the changes in motif sensitivity described in Supplementary Note 3.3. Accordingly, we see that rankings are more affected by merging with relaxed parameters.

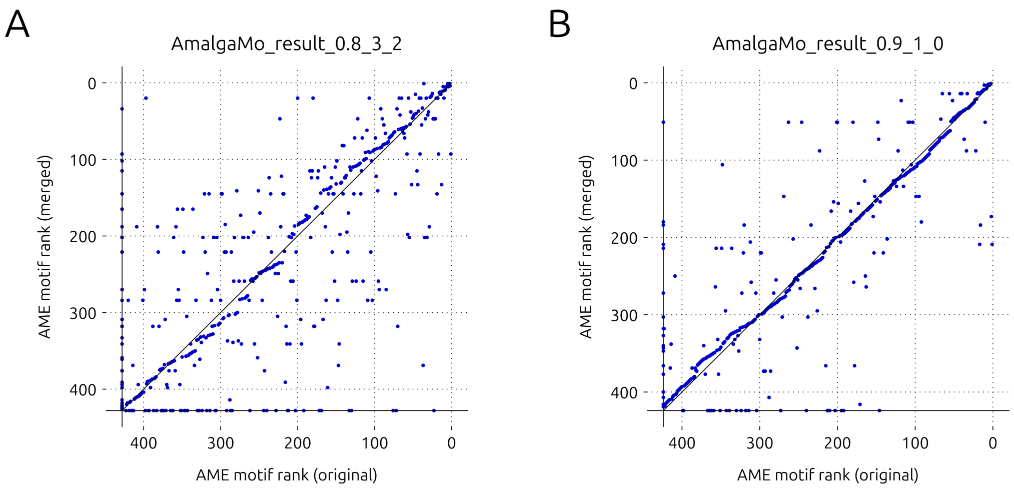

**Supplementary Figure SN5.9:** AME motif enrichment ranking before and after merging. A. Motif set obtained using the most relaxed parameter set. B. Motif set obtained using the most strict parameter set.

### Supplementary Tables

**Supplementary Table S1:** Datasets used in this study. hESC: human embryonic stem cell line. OE: overexpression. CVID: common variable immunodeficiency.

| reference | SRA accession | biological sample | condition | # replicates |
|---|---|---|---|---|
| Pahl et al. (2024) | PRJNA960640 | human naïve CD4 T cells | unstimulated<br>stimulated (8h) | 3<br>3 |
| Li et al. (2019) | PRJNA422918 | differentiated H9 hESCs | TFAP2C OE off<br>TFAP2C OE on | 2<br>2 |
| Mandal et al. (2018) | PRJNA400057 | human EBV-transformed B cells | healthy<br>CVID | 5<br>4 |
